# The spatiotemporal neural dynamics of object recognition for natural images and line drawings

**DOI:** 10.1101/2022.08.12.503484

**Authors:** Johannes J.D. Singer, Radoslaw M. Cichy, Martin N. Hebart

**Author notes:** These authors contributed equally (Radoslaw M. Cichy, Martin N. Hebart).

## Abstract

1.

Drawings offer a simple and efficient way to communicate meaning. While line drawings capture only coarsely how objects look in reality, we still perceive them as resembling real-world objects. Previous work has shown that this perceived similarity is mirrored by shared neural representations for drawings and natural images, which suggests that similar mechanisms underlie the recognition of both. However, other work has proposed that representations of drawings and natural images become similar only after substantial processing has taken place, suggesting distinct mechanisms. To arbitrate between those alternatives, we measured brain responses resolved in space and time using fMRI and MEG, respectively, while human participants (female and male) viewed images of objects depicted as photographs, line drawings, or sketch-like drawings. Using multivariate decoding, we demonstrate that object category information emerged similarly fast and across overlapping regions in occipital, ventral-temporal and posterior parietal cortex for all types of depiction, yet with smaller effects at higher levels of visual abstraction. In addition, cross-decoding between depiction types revealed strong generalization of object category information from early processing stages on. Finally, by combining fMRI and MEG data using representational similarity analysis, we found that visual information traversed similar processing stages for all types of depiction, yet with an overall stronger representation for photographs. Together our results demonstrate broad commonalities in the neural dynamics of object recognition across types of depiction, thus providing clear evidence for shared neural mechanisms underlying recognition of natural object images and abstract drawings.

**Significance Statement:** When we see a line drawing, we effortlessly recognize it as an object in the world despite its simple and abstract style. Here we asked to what extent this correspondence in perception is reflected in the brain. To answer this question, we measured how neural processing of objects depicted as photographs and line drawings with varying levels of detail (from natural images to abstract line drawings) evolves over space and time. We find broad commonalities in the spatiotemporal dynamics and the neural representations underlying the perception of photographs and even abstract drawings. These results indicate a shared basic mechanism supporting recognition of drawings and natural images.

## 3. Introduction

Line drawings are universal in human culture and provide a simple and efficient tool for visualization. With just a few strokes we can depict the things that we encounter in everyday life in a way that is easily recognizable by others. Line drawings of objects can be recognized without any previous experience (Kennedy & Ross, 1975), by infants only a few months after birth (DeLoache et al., 1979), and across a large variation of styles and levels of detail of the drawing (Eitz et al., 2012). This ease of recognition raises the question as to how line drawings convey meaning so efficiently.

One possible explanation for our ability to recognize line drawings efficiently is that they resemble natural object images in terms of some core visual features that are central to object recognition (Fan et al., 2018). It has been suggested that these visual features correspond to the edges of an image (Biederman & Ju, 1988). Considering the architecture of visual cortex, it has been proposed that lines in drawings drive early visual brain areas in a similar fashion to edges in natural images and therefore lead to a similar representation of objects in the brain (Sayim & Cavanagh, 2011). This notion is supported by work demonstrating that the recognition of object drawings engages the same brain regions as photographs (Ishai et al., 2000; Kourtzi & Kanwisher, 2000) and that early and high-level visual brain regions similarly represent category information for drawings and natural object images (Haxby et al., 2001; Spiridon & Kanwisher, 2002). While these results indicate that drawings and natural object images share a representational format in some visually responsive brain regions, the exact spatial extent, the temporal dynamics, and the spatiotemporal evolution of the similarities in processing of natural object images and drawings remain largely unknown.

An alternative explanation for the recognition of drawings is that the visual information retained in line drawings is too abstract and therefore insufficient to drive visual recognition mechanisms tuned to natural images. According to this view, additional processing steps are required to refine the representation of drawings, making it more similar to the representation of natural images over time. For scenes, there is evidence suggesting that similarities in processing of natural scene images and scene drawings become progressively stronger with the depth of visual processing (Walther et al., 2011) or even emerge only late in time (Lowe et al., 2018). In addition, previous results in support of a shared representational format for natural object images and drawings (Haxby et al., 2001; Spiridon & Kanwisher, 2002) used fMRI alone, making it impossible to infer whether the effects were driven by the same or distinct underlying temporal dynamics for drawings and natural images. This leaves open whether drawings and natural object images are similarly processed from early on or whether the shared representational format is a result of additional processing steps required for drawings.

To provide evidence in favor or against these explanations, here we resolved the similarities and differences in processing of natural object images and drawings across space and time. To this end, we measured fMRI and MEG in two sessions while participants viewed object images depicted across three levels of visual abstraction: colored photographs, detailed black-and-white line drawings, and abstract sketch-like drawings. Using spatially and temporally-resolved multivariate decoding and representational similarity analysis (RSA, Kriegeskorte et al., 2008), we provide clear evidence in favor of common representational dynamics for objects across levels of visual abstraction in visual cortex. These results elucidate the representational nature of drawings in visual cortex and suggest common neural mechanisms for object recognition across levels of visual abstraction.

## 4. Materials and Methods

### 4.1. Participants

Thirty-one healthy adults with normal or corrected-to-normal vision took part in the study and provided their written informed consent before participating. In total, we excluded 8 participants from the analysis of the fMRI data and 9 from the analysis of the MEG data. We based the exclusion on withdrawn participation (one participant, both fMRI and MEG sessions), low alertness (>20% missed catch trials, see Experimental Task Paradigm, 2 fMRI sessions and 5 MEG sessions), missing data (no structural image, one fMRI session), noisy data (>1% outlier volumes in framewise intensity difference / excessive head motion, 4 fMRI sessions), and excessive eye movements on the stimulus (>5% of experimental trials, see section on eye movement recording and analysis, affecting 3 MEG sessions). Hence, for the fMRI analyses, we included data of 23 participants (mean age=29.22, SD=3.97, 13 female, 10 male), while for the MEG analyses, we included 22 partly overlapping participants (mean age=28.91, SD=4.02, 10 female, 12 male, 17 overlapping with fMRI analysis). Please note that post-hoc analyses including all the subjects into the analysis for whom data was available did not qualitatively change the pattern of results, demonstrating that exclusion criteria did not alter the overall pattern of results. The study was approved by the local ethics committee of the University Medical Center Leipzig (012/20-ek) in accordance with the declaration of Helsinki, and participants were reimbursed for their participation.

### 4.2. Experimental stimuli

We used object images of the same 48 categories in three different types of depiction (144 stimuli in total), each representing one level of visual abstraction (Fig. 1a). 24 of these object categories were natural objects (e.g. animals and plants), while the other 24 were man-made (e.g. food, tools and vehicles). For each category and type of depiction there was one exemplar. For the first type of depiction (“photos”), we used colored photographs of objects, cropped from their background. For the second type of depiction (“drawings”), we asked an artist to draw black and white line drawings based on the photos with a high level of detail. In these drawings, color and some texture features were abstracted while retaining most of the contours of the objects. Finally, in the third type of depiction (“sketches”), the artist was instructed to draw line drawings of the photos in a highly abstracted way. In comparison to the drawings and photos, the sketches distorted the contours and the size of some parts of the objects, and texture information was reduced to a minimum.

**Figure 1.**
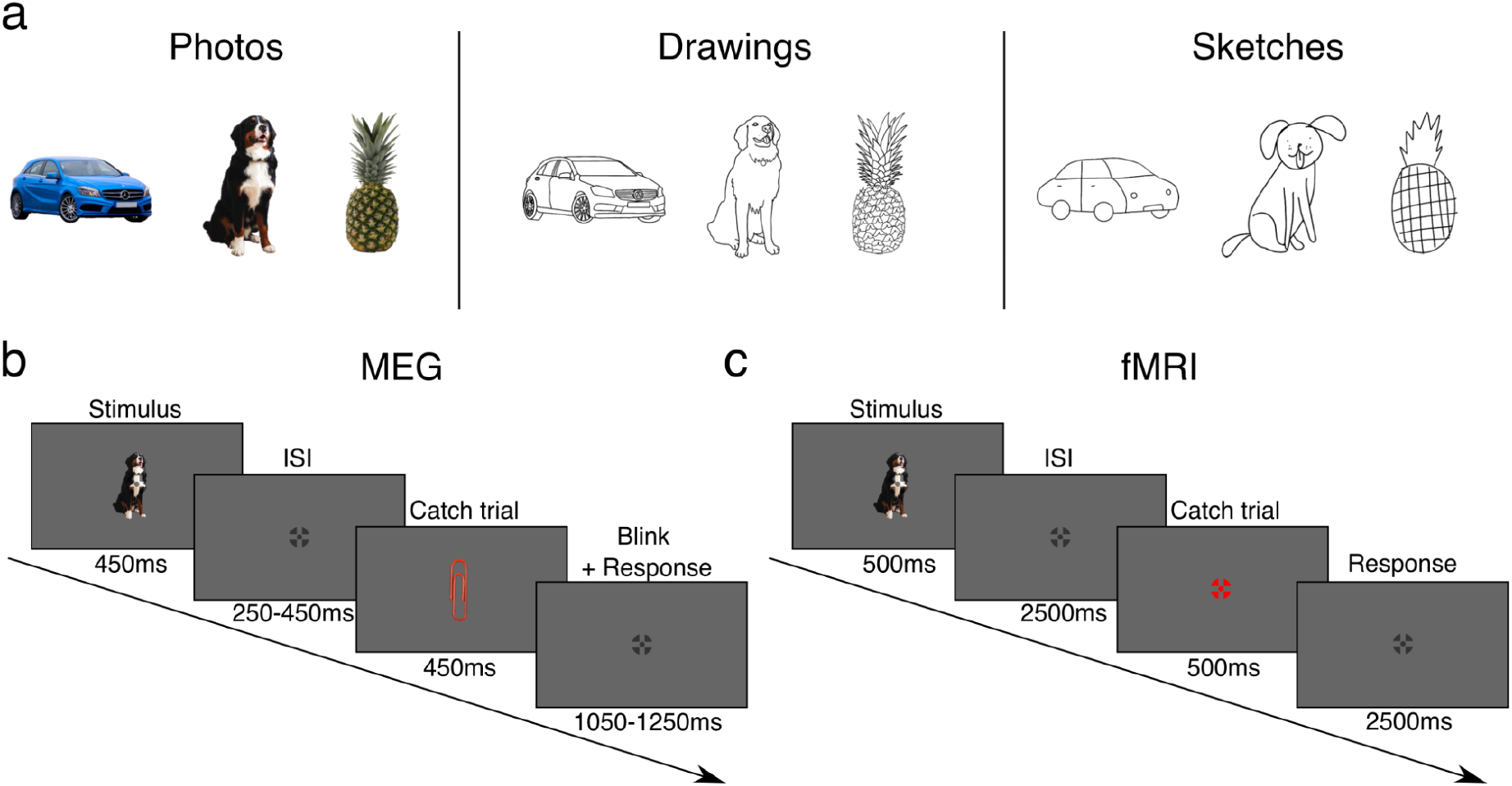
Stimuli and experimental paradigm. **a) Stimulus set used in the experiment.** We used images of the same 48 object categories in three types of depiction (144 stimuli in total). Objects were depicted as either photos, drawings, or sketches, with each type of depiction reflecting a different level of visual abstraction. **b) MEG paradigm.** In the MEG experiment, participants viewed sequences of object images in random order while fixating on a central fixation cross. Their task was to respond to rare catch trials by pressing a button and blinking. **c) fMRI paradigm.** Analogous to the MEG experiment, in the fMRI experiment participants viewed sequences of object images in random order while fixating on the central fixation cross. Object sequences were interspersed with catch trials in which participants were instructed to respond with a button press. Stimulus presentation timing and ISIs were adjusted according to the modality-specific requirements.

#### 4.2.1. Quantitative validation of the experimental stimuli

To be able to meaningfully compare object recognition for photographs and drawings at different levels of visual abstraction we reasoned that our stimulus set is required to suffice two main criteria: stimuli in the three types of depiction should (1) differ in terms of their low-level visual features, reflecting a difference in the degree of visual abstraction and (2) be perceived similarly at a conceptual level by human participants.

First, to quantitatively validate that the stimuli in the three types of depiction differ in their level of visual abstraction, we extracted low-level visual features for them using the deep convolutional neural network VGG16 (Simonyan & Zisserman, 2015). The network is widely used and prominent for its appearance at the ImageNet Large Scale Visual Recognition Challenge (Russakovsky et al., 2015) in 2014, where it reached a top-5 test accuracy of 92.7% on the ImageNet dataset. VGG16 contains five convolutional blocks, each composed of a series of convolutional layers, followed by a max pooling and a ReLU layer. After the convolutional layers there are three fully connected layers. The last fully connected layer outputs class probability values for all of the 1,000 classes in the ImageNet dataset (Deng et al., 2009) after applying a softmax activation function. We used VGG16 as it has not only achieved good performance in image recognition tasks but also repeatedly has been shown to learn representations that resemble visual object representations in the human brain (Güçlü & Gerven, 2015; Schrimpf et al., 2020; Storrs et al., 2021). As an approximation for low-level visual feature representations, we extracted network activations from pooling layer 2 in response to our object images (Bankson et al., 2018; Greene & Hansen, 2020; Reddy et al., 2021; Xie et al., 2020). For feeding the object images through the network, the objects were put on a square gray background and resized to 224 x 224 pixels. Next, we computed representational dissimilarity matrices (RDMs, Kriegeskorte et al., 2008) by correlating all activations in a given type of depiction with each other and computing pairwise distances by using 1-Pearson correlation as a distance measure. This yielded one low-level visual RDM for each type of depiction. We finally compared these RDMs by correlating their lower triangular parts to each other using Pearson correlation. This resulted in one correlation value for a given comparison between two types of depiction (e.g. photo-drawing), reflecting the degree of low-level feature similarity of the stimuli.

To ensure that human participants perceive the stimuli in the different types of depiction similarly at a conceptual level, we used data from a previous study (Singer et al., 2022) in which workers on Amazon Mechanical Turk had performed a triplet odd-one out task (Hebart et al., 2020) on the same stimuli as used here. In this task participants were instructed to find the odd-one out in triplets of object images belonging to the same type of depiction. Based on these triplet judgments we constructed human perceptual similarity matrices for each type of depiction separately, describing the representational object space based on human behavior. Subsequently, we correlated the lower triangular parts of the similarity matrices from the different types of depiction using Pearson correlation, yielding a measure of perceptual similarity between all types of depiction.

### 4.3. Experimental design and procedure

All participants first completed one fMRI experiment, followed by an MEG experiment on a separate day, which took place on average 30.57 days after the first experiment (range 7-85). Before the fMRI experiment, participants were familiarized with the stimuli used in both experiments. This was done to ensure that every participant was able to recognize the objects depicted in all of the images in order to rule out the possibility of differences between types of depiction based on the recognizability of the images.

#### 4.3.1. Experimental paradigm

During both experiments (fMRI, MEG), subjects were presented with images of the same object categories in three types of depiction (photos, drawings, sketches). Depiction types were not mixed within runs but presented in separate runs to avoid carry-over effects of consecutive presentation of the same object in different types of depiction. Participants were instructed to maintain fixation at the center of the screen indicated by a fixation cross during the whole experiment (Fig. 1b-c).

Stimuli were presented at the center of the screen overlaid with a semitransparent crosshair fixation cross (Thaler et al., 2013), which subtended 0.63° in the fMRI experiment and 0.5° of visual angle in the MEG experiment. The individual stimulus size was manually adjusted before the experiment such that the area an object image occupied on the screen was approximately equal for all object images. Hence, the stimulus size could vary across object images, and one object image subtended on average 4.34° (Range = [2.97°, 5.85°]) in the fMRI experiment and 6.15° of visual angle (Range = [4.21°, 8.25°]) in the MEG experiment.

Stimulus presentation timings were adjusted to the specifics of the imaging modality. In the MRI experiment, each stimulus was presented for 500ms followed by an interstimulus interval (ISI) of 2500ms (total trial duration 3s). In the MEG experiment, each stimulus was presented for 450ms followed by an ISI which was randomly sampled from a range of values between 250ms to 450ms in steps of 50ms to reduce effects of phase synchronization (average total trial duration 800ms).

In both experiments, stimulus presentations were interleaved with catch trials in which participants were instructed to respond to a given stimulus, in order to keep the subjects alert. In the MRI experiment, participants were instructed to respond with a button press when the fixation cross turned red. In the MEG experiment, they were instructed to respond to a paperclip stimulus (which was presented in the type of depiction of the corresponding run e.g., as a drawing) and to blink, in order to reduce blinking artifacts during the experimental trials. In the MRI experiment, the ISI for catch trials was equal to the ISI of experimental trials (total trial duration 3s). Catch trials in the MEG experiment were followed by a longer ISI (range of values between 1,050ms and 1,250ms in steps of 50ms) to give participants time to respond and for the MEG signal to return back to baseline after the blink (average total trial duration 1600ms).

In a given run, each object image of a given type of depiction was presented twice in the MRI and eight times in the MEG experiment. Stimulus presentation order was randomized while prohibiting immediate stimulus repetition. Catch trials accounted for 20% of the trials in both experiments and were presented after every 4th to 6th object image presentation. In total, each participant completed 12 runs in the MRI experiment (4 from each condition, randomized in order, total run duration 6min 16.5s) and 9 runs in the MEG experiment (3 from each condition, randomized in order, total run duration 7min 44.8s), resulting in 8 stimulus presentations per image and condition across runs in the MRI experiment and 24 stimulus presentations per image and condition across runs in the MEG experiment.

#### 4.3.2. Functional localizer task

Before the experimental task in the fMRI experiment, participants underwent one functional localizer run independent from the experimental runs, which was later used for defining regions of interest (ROIs). Subjects were presented with either fully visible object images (objects), scrambled object images (scrambled), or a fixation cross. Participants were instructed to fixate on the fixation cross and to respond with a button press if the same object was presented in two consecutive trials. Objects and scrambled objects were presented at the center of the screen for a duration of 400ms, followed by a presentation of a fixation cross for 350ms. Both types of images were presented in blocks of 15s each and interleaved with blocks of 7.5s of fixation. The localizer run comprised 12 blocks of fixation and 12 blocks of both objects and scrambled objects with a total run duration of 7min 45s.

### 4.4. fMRI acquisition, preprocessing and univariate analysis

#### 4.4.1. fMRI acquisition

We recorded fMRI data on a Siemens Magnetom Prisma Fit 3T system (Siemens, Erlangen, Germany) using a 32-channel head coil. Functional images were acquired using a multiband 3 sequence (TR=1.5s, TE=33.2ms, in-plane resolution: 2.49×2.49mm, matrix size=82×82, FOV=204mm, flip angle=70°, 57 slices, slice thickness=2.5mm) with whole brain coverage. Existing T1-weighted structural images obtained in previous studies were used that varied in exact sequence parameters (MPRAGE, voxel size = 1mm^3^).

#### 4.4.2. fMRI preprocessing

All preprocessing and univariate analyses of the fMRI data were conducted using SPM12 (https://www_l.ion.ucl.ac.uk/spm/) and custom scripts in Matlab R2021a (www.mathworks.com).

First, we screened functional data for outliers in image intensity difference and head motion. To this end, we carried out initial realignment and computed the difference in image intensity of each functional volume and its subsequent volume for each brain slice, excluding the eyeballs. Next, to determine outlier volumes, we scaled the differences between functional images relative to the overall mean of differences across all functional images. We excluded subjects for whom more than 1% of volumes showed a more than 30-fold increase in image intensity difference or a displacement of more than 0.5mm in any direction. For all other subjects, we removed and then linearly interpolated the images that exceeded the criteria.

Following outlier removal, functional images were realigned to the first image of the run, slice-time corrected, and coregistered to the anatomical image. The functional images of the localizer task were smoothed with a Gaussian kernel (FWHM = 5mm) while the images from the experimental runs were not smoothed.

Further, we estimated noise components for the functional images of the experimental runs by using the aCompCor method (Behzadi et al., 2007) implemented in the TAPAS PhysIO toolbox (Kasper et al., 2017). To this end, tissue-probability maps for the gray matter, white matter, and cerebrospinal fluid (CSF) were estimated based on the structural image of a participant, and noise components were extracted based on the tissue-probability maps of the white matter and CSF in combination with the fMRI time series.

#### 4.4.3. fMRI univariate analysis

We modeled the fMRI responses to each object image in a given run with a general linear model (GLM). The onsets and durations of each object image were entered as regressors into the model and were convolved with a hemodynamic response function (HRF) resulting in 48 regressors for the experimental conditions in each run. As nuisance regressors, we included the noise components extracted from the white matter and CSF maps as well as the movement parameters and their first and second order derivatives. We repeated this GLM approach 20 times, each time convolving with a different HRF obtained from an openly available library of HRFs (https://github.com/kendrickkay/GLMsingle) which was derived from a large fMRI dataset of participants viewing natural scenes (Allen et al., 2022). After fitting the GLMs, for each voxel we extracted the beta values for the object image regressors from the GLM with the HRF that had resulted in the minimum mean residual for that given voxel. Since the true HRF is variable across subjects, tasks and even brain regions (Polimeni & Lewis, 2021) this approach allows a closer approximation of the true HRF in comparison to using the canonical HRF while it does not lead to positively biased statistics at the group level. This procedure yielded 48 beta maps (one for each object category) for each run and participant. For later searchlight analyses, we normalized these beta maps to the MNI template brain.

The fMRI responses for the localizer experiment were modeled in a separate GLM, with the onsets and durations of the blocks of objects and scrambled objects convolved with the canonical HRF as regressors. Only movement parameters were included as nuisance regressors in this GLM. From the resulting beta estimates we computed three contrasts. The first contrast was used to localize activity in early visual brain areas and was defined as scrambled > objects. The second contrast was used to localize activity in object-selective cortex and was defined as objects > scrambled. The third contrast was used to localize activity in posterior parietal cortex and was defined as objects+scrambled > baseline. This way, we obtained three *t*-maps for the three contrasts for each participant.

#### 4.4.4. Region-of-interest definition

We focused on regions in early visual cortex (EVC) i.e., V1, V2, V3, the lateral occipital complex (LOC), comprising object-selective regions LO and pFs in the ventral stream, and on the posterior intraparietal sulcus (pIPS), comprising the regions IPS0 and IPS1 in the dorsal stream.

To define EVC, we first transformed the subject-specific *t*-maps from the scrambled > objects contrast from the localizer GLM into MNI-space. Based on these transformed *t*-maps we computed a contrast comparing the group-level activation against zero, which resulted in one *t*-map across subjects. We then thresholded this *t*-map at the *p*<0.001 level and calculated the overlap between the thresholded *t*-map and the combined anatomical definition of V1, V2 and V3 from the Glasser Brain Atlas (Glasser et al., 2016). Finally, we transformed this overlap image back into the native subject space for each subject resulting in subject-specific EVC masks. Please note that a more fine-grained definition of the ROIs V1, V2, V3, and V4 based on the Wang et al. (2015) atlas led to qualitatively similar results as the EVC definition.

To define object-selective cortex, we manually identified the peaks in the subject-specific *t*-maps of the objects > scrambled contrast from the localizer GLM which corresponded anatomically to LO and pFS. We then defined spheres with a radius of 6 voxels around both peaks, including only those voxels in the spheres that had *t*-values corresponding to *p*<0.0001. This resulted in one ROI mask for LO and pFS, respectively. Initial exploratory analyses revealed that LO and pFS yielded highly comparable results. Therefore, we merged the two ROI masks into one combined LOC mask. This resulted in one object-selective cortex mask for each subject.

To define pIPS, we first combined the probability masks for IPS0 and IPS1 from the Wang et al. (2015) atlas and then thresholded this combined IPS0-1 mask at a value of 20%. Next, we transformed the combined pIPS mask into the individual subject space. Finally, we computed the overlap between the individual pIPS mask and the subject-specific *t*-map of the contrast from the localizer GLM comparing all objects and scrambled objects against baseline, thresholded at *p*<0.0001. This procedure resulted in one pIPS ROI mask for each subject. In case the EVC, object-selective cortex, or pIPS masks overlapped in a given subject, the overlapping voxels were discarded from all masks.

### 4.5. MEG acquisition and preprocessing

#### 4.5.1. MEG acquisition

Before the MEG measurement started, participants’ head shape was digitized using a Polhemus FASTRAK device. Additionally, five coils were placed on the head of the participant which were later used to track the head position inside the MEG. During the experiment that took place inside a magnetically shielded room, we recorded neuromagnetic signals using a 306-channel NeuroMag VectorView MEG system (Elekta, Stockholm) with a sampling rate of 1,000Hz and an online filter between 0 and 330Hz.

#### 4.5.2. MEG preprocessing

To remove external noise and correct for head movements during the MEG measurement, we applied temporal signal space separation (Taulu & Simola, 2006) and movement correction to the MEG data using the Maxfilter software (Elekta, Stockholm). All further preprocessing steps were implemented in Matlab R2021a (www.mathworks.com), using the utilities of the Fieldtrip toolbox (Oostenveld et al., 2011) and custom scripts.

First, Independent Component Analysis (ICA) was applied to the combined data from all blocks to identify components corresponding to eye movements, blinks, or heartbeat. The resulting ICA components were manually inspected in combination with their topographies and time courses, and only those components that could be clearly attributed to eye movements, blinks, or heartbeat were removed from the data. Using this procedure, for a given subject, we removed an average of 1.73 components (SD = 0.69). Please note that the removal of eye movement and blink-related independent components is only meant to clean the data from noise related to these components and is unrelated to the exclusion criteria based on eye movements on the stimulus. Next, the data were filtered with a 0.5Hz high pass filter and a 40Hz low pass filter and segmented into trials starting 100ms prior to the onset of a given stimulus and ending 1,001ms after the stimulus presentation. Importantly, triggers indicating the beginning of the stimulus presentation were adjusted to match the exact time of the onset of a given image presentation by aligning them to the onset of the response of an optical sensor attached to the projection monitor in the MEG. Following this step, data were baseline corrected with respect to the time period −100ms to 0ms relative to stimulus onset and downsampled to 100Hz to speed up later multivariate analyses. Finally, multivariate noise normalization was applied to the data (following general guidelines for multivariate pattern analysis of M/EEG data (Guggenmos et al., 2018)). In sum, this procedure resulted in trials of 111 timepoints across 306 channels for every participant.

### 4.6. Eye movement recording and analysis

During the MEG experiment, eye movements of the subject were recorded using an SR Research EyeLink 1000 system (SR Research Ltd). These data were only acquired for the purpose of identifying subjects that made a significant amount of eye movements on the presented stimulus, which might be informative about the stimulus identity and could therefore bias results of multivariate pattern analysis (Mostert et al., 2018; Thielen et al., 2019). No reliable eye movement data could be acquired for 4 subjects, so they were excluded from further eye movement data analyses.

First, the data were filtered with a 0.1Hz high pass filter to remove slow drifts, followed by segmentation into epochs beginning 100ms prior to and ending 500ms after stimulus onset. Second, we removed epochs that contained estimated eye movements with an amplitude greater than 3° of visual angle based on the assumption that these movements could not have fallen on the presented stimulus and thus could not constitute an eye movement on the stimulus but rather must reflect noise or occasional non-informative eye movements beyond the stimulus. Finally, we discarded the pupil diameter channel from the data and retained only the horizontal and vertical position channels for further analyses.

As an index for an eye movement on the stimulus, we detected saccades and microsaccades in the extracted clean epochs by using the microsaccade detection algorithm by Engbert & Kliegl (2003). Subsequently, we computed the amplitude of movement in a given detected micro-saccade and labeled only the micro-saccades with an amplitude greater than 1.5° of visual angle as eye movements on the stimulus, given that any smaller eye movements would be hard to distinguish from noise. Finally, we computed the ratio of trials containing eye movements on the stimulus to all experimental trials (excluding catch trials) to determine how many experimental trials were contaminated by eye movements on the stimulus for a given subject. Based on this estimate, we excluded three participants from the MEG analysis because they showed such eye movements in more than 5% of the remaining experimental trials.

### 4.7. Multivariate decoding of object category information

We used multivariate decoding on the preprocessed fMRI voxel patterns and MEG channel patterns to determine where and when the category information of a presented object can be read out from brain activity. To this end, separately for every type of depiction, we trained and tested linear Support Vector Machine (SVM) classifiers (Chang & Lin, 2011) to distinguish between the responses to two given objects for every possible combination of objects, resulting in one accuracy value for every pair of objects (50% chance level). Subsequently, we averaged all pairwise accuracies to obtain a measure of overall object discriminability. This procedure was repeated across ROIs or searchlights for the fMRI data and across time points for the MEG data. All decoding analyses were performed separately for every participant.

#### 4.7.1. Spatially-resolved multivariate fMRI decoding

To ask where in the brain category information can be read out from fMRI voxel activity patterns, we used both an ROI-based and a spatially unbiased searchlight procedure (Haynes et al., 2007; Kriegeskorte et al., 2006).

For the ROI-based procedure, we arranged the beta values from the voxels in a given ROI into pattern vectors for each object category and run. We then evaluated classifiers using a leave-one-out cross-validation procedure, training on the pattern vectors from three runs and testing on the pattern vector from the remaining run. We repeated this procedure until every pattern vector had been used once for testing (see Fig. 2a for visualization of the approach). This resulted in decoding accuracies for every ROI, each type of depiction, and each participant.

**Figure 2.**
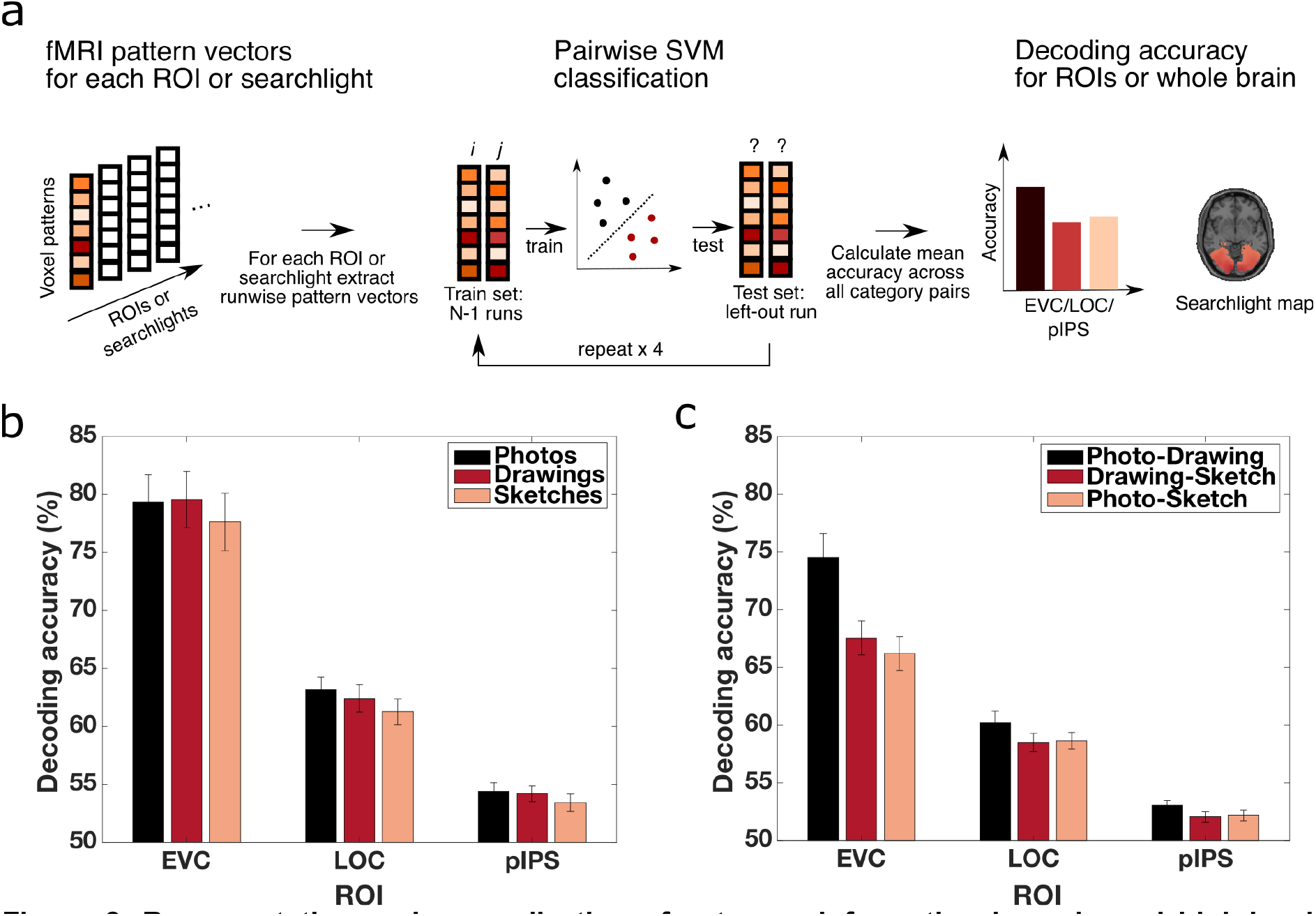
Representation and generalization of category information in early and high-level visual cortex at different levels of visual abstraction. **a) Spatially resolved decoding procedure.** We trained SVM classifiers on the voxel activity patterns of a given ROI or searchlight to classify if a given pattern belonged to object category *i* or *j* for all possible pairs of objects using a leave-one-out cross-validation framework. Subsequently, we averaged the pairwise decoding accuracies for all object pairs, resulting in decoding accuracies across ROIs or searchlights for each participant and type of depiction separately. **b) Category decoding accuracies in early and high-level visual cortex across levels of visual abstraction.** We found above chance decoding accuracies for all types of depiction in EVC, LOC and pIPS. There were no significant differences in decoding accuracies between types of depiction in any of the ROIs. **c) Cross-decoding accuracies between types of depiction across ROIs.** We found significant cross-decoding accuracies between all types of depictions in EVC, LOC as well as pIPS. Error bars reflect the standard error of the mean across participants.

For the spatially unbiased searchlight analysis, we defined a sphere with a radius of 4 voxels around a given voxel and formed pattern vectors based on all the beta values within this sphere. Analogous to the ROI-based procedure, we then evaluated classifiers using a leave-one-out cross-validation procedure. This evaluation procedure was iterated over all possible searchlights, yielding accuracy values across the whole brain for each type of depiction and each participant separately. The resulting searchlight maps were subsequently smoothed with a Gaussian kernel (FWHM = 5mm).

#### 4.7.2. Temporally-resolved multivariate MEG decoding

For the temporally-resolved decoding analyses, we arranged the preprocessed MEG data into pattern vectors containing the MEG data across channels for every object category, trial and time point. Subsequently, to improve the signal-to-noise ratio, we averaged data from two trials of the same object category into one supertrial, resulting in twelve supertrials per object category and time point. We then evaluated SVM classifiers using a leave-one-out cross-validation framework, training the classifiers on eleven supertrials and testing on the left out supertrial and repeating this procedure until every supertrial had been used once for testing (see Fig. 5a for visualization of the approach). To increase the robustness of the results, we repeated the whole cross-validation procedure and the averaging of trials into supertrials five times while randomizing the assignment from trials to supertrials. Accuracies were subsequently averaged across repetitions. This procedure was repeated for every time point and for each type of depiction separately, which resulted in object decoding time courses for every type of depiction and every participant.

#### 4.7.3. fMRI and MEG cross-decoding of category information between types of depiction

To determine where and when object category information generalizes between types of depictions, we used cross-decoding. This approach was analogous to the regular decoding procedure, but instead of training and testing on data from the same type of depiction, we trained a classifier on data from one type of depiction (e.g., photos) and tested on data from another type (e.g., drawings). We carried out cross-decoding for three types of comparisons: photo-drawing, photo-sketch and drawing-sketch. Further, we computed the cross-decoding accuracies for both train-test directions and averaged the accuracies subsequently. This way, data from both types of depiction was used once for training and once for testing the classifier. Analogous to the regular decoding procedure, we repeated this procedure across ROIs and searchlights for fMRI and across time points for MEG data, resulting in cross-decoding accuracies across space and time for the three comparisons and for each participant separately.

#### 4.7.4. MEG temporal generalization analysis

To investigate at which points in time the object category MEG pattern information generalized to other points in time, we used the temporal generalization method (King & Dehaene, 2014). For a given time point, we trained a classifier analogous to the temporally-resolved decoding procedure. To determine the generalization of this classifier across time, we tested the classifier on patterns not only at the matching time point but at all timepoints. Then, we repeated this training-generalization approach for every time point, yielding a time × time temporal generalization matrix of decoding accuracies for each type of depiction and each participant.

### 4.8. RSA-based MEG-fMRI fusion

For combining the information about visual processing in the spatial dimension from fMRI data with the temporal dimension from MEG data, we used RSA-based MEG-fMRI fusion (Cichy et al., 2014; Cichy & Oliva, 2020; Hebart et al., 2018; see Fig.8a for a visualization of the approach). The basic idea behind RSA is to characterize the representational space in a given measurement component (e.g. an fMRI ROI) with an RDM. An RDM describes the representational space in terms of pairwise distances between responses to all of the conditions of interest, thereby abstracting from the activity patterns of measurement channels (e.g. fMRI voxels or MEG sensors). RDMs can be obtained e.g. across different regions in the brain or points in time and can subsequently be compared by correlating them. If two RDMs exhibit a positive correlation, it is assumed that the underlying representational geometry is similar. Following this rationale, we computed RDMs for each time point, ROI, type of depiction and each subject separately. For this, we first averaged all run-wise fMRI or trial-wise MEG pattern vectors for a given object category extracted at different ROIs or time points. Subsequently, we computed the pairwise dissimilarities between pattern vectors as 1 - Pearson correlation and stored these dissimilarities in one RDM for a given ROI or time point. Then, we correlated the lower triangular parts of the ROI-specific and temporally-resolved RDMs with each other using Pearson correlation, resulting in MEG-fMRI fusion time courses for each ROI, each type of depiction and each participant separately.

### 4.9. Statistical analyses

To assess the statistical significance of the decoding accuracies as well as RDM correlations, we used non-parametric sign-permutation tests (Nichols & Holmes, 2002). To this end, we obtained null distributions by randomly permuting the sign of the results at the participant level a total number of 10,000 times. Based on these null distributions, we obtained *p*-values for the empirical results and thresholded these *p-* values at the *p*<0.001 level. *P*-values obtained for decoding accuracies were based on one-sided tests, while *p*-values for RDM-correlations as well as differences of decoding accuracies were based on two-sided tests. Uncorrected *p*-values were only used for inference when testing decoding accuracies against chance in individual ROIs since results for photos, drawings, and sketches were treated as testing separate hypotheses. However, when testing for pairwise differences between conditions (i.e. photo vs. drawing, photo vs. sketch, drawing vs. sketch) or when testing cross-decoding accuracies for multiple combinations of depiction types (i.e. photo-drawing, photo-sketch, drawing-sketch) against chance within a given ROI, we corrected the *p*-values with the Benjamini-Hochberg FDR correction (Benjamini & Hochberg, 1995).

For statistical tests across voxels or time involving a large number of multiple comparisons, we applied cluster correction to control the alpha-error rate (Maris & Oostenveld, 2007). The data points that exceeded the *p*<0.001 threshold were clustered based on temporal or spatial adjacency, and the maximum cluster size was computed for each permutation. This way, we obtained a null-distribution of the maximum cluster size statistic. Finally, the clusters in the empirical results were then thresholded based on the null-distribution of the maximum cluster size statistic at the *p*<0.05 level. To correct for multiple tests of significance of pairwise differences between conditions (e.g., photo vs. drawing, photo vs. sketch, drawing vs. sketch) or for testing cross-decoding accuracies for multiple combinations of depiction types (i.e. photo-drawing, photo-sketch, drawing-sketch), we obtained the cluster-size statistic which corresponded to the given statistical threshold (*p*<0.05) for all of the multiple tests and used the maximum cluster-size statistic computed across tests as the threshold for all clusters from all tests.

In order to estimate confidence intervals for the decoding accuracy and RDM correlation peak latencies, we used a bootstrapping procedure. For this, we randomly sampled participant specific time series with replacement for a total number of 100,000 times. Next, we averaged the results across participants for every bootstrap sample and then estimated the peak latency by finding the maximum of the average time series. Based on the mean and standard deviation of the resulting distribution of peak latencies we computed the 95% confidence intervals of the peak latency.

For comparing decoding accuracy and RDM correlation peak latencies we used a bootstrapping procedure analogous to the approach described above. However, instead of estimating confidence intervals of peak latencies of one condition we estimated the confidence intervals of the difference between conditions by subtracting the peak latencies for two given conditions estimated for each bootstrap sample. This yielded a distribution of peak latency differences from which we obtained the 95% confidence interval of the difference. We regarded a given difference between peak latencies as significant if the confidence interval of the difference did not include zero.

Finally, to test for the statistical equivalence of decoding accuracy or RDM correlation peak latencies we used a two one-sided tests procedure (TOST) (Lakens, 2017).

### 4.10. Data and code availability

All results of the decoding and RSA analyses are publicly available via https://osf.io/vsc6y/ along with preprocessed fMRI and MEG data from an exemplary subject. The raw MEG and fMRI data can be accessed on OpenNeuro via https://openneuro.org/datasets/ds004330 and https://openneuro.org/datasets/ds004331. Code to reproduce the results and figures in the paper is provided via https://github.com/Singerjohannes/object_drawing_dynamics.

## 5. Results

### 5.1. Natural object images and line drawings differ in low-level visual features but are perceived similarly

To ensure that our stimulus set is well suited for comparing object recognition across different levels of visual abstraction, we aimed to quantitatively validate that objects are perceived similarly by human subjects at a conceptual level despite differences at the visual level. As a proxy for low-level visual features, we first extracted features from pooling layer 2 of the deep convolutional neural network VGG16 (Simonyan & Zisserman, 2015) for all of the object images, in line with previous work (Bankson et al., 2018; Greene & Hansen, 2020; Reddy et al., 2021; Xie et al., 2020). We then computed RDMs based on the extracted features separately for the different types of depiction and correlated the lower triangular parts of the RDMs between types of depiction. As expected, photos and drawings showed the highest RDM correlation (*r*=0.79) while the correlation for photos and sketches (*r*=0.41) as well as the correlation between drawings and sketches (*r*=0.45) were lower. Next, to confirm that human subjects perceive the object images in the different types of depiction similarly at a conceptual level, we used previously acquired data (Singer et al., 2022) where workers on Amazon Mechanical Turk indicated which of three object images they thought was the odd-one out (Hebart et al., 2020). These triplet judgments were used to construct perceptual similarity matrices for each type of depiction separately, which we subsequently correlated to each other to estimate their representational similarity. As expected, human subjects perceived all types of depictions highly similarly (all pairwise correlations *r*=0.97). In sum, these analyses quantitatively confirm that while there is a gradual difference in low-level visual features across the three types of depiction reflecting the degree of visual abstraction, there is also a correspondence in how human participants perceive these images at a conceptual level.

### 5.2. Object category information can be decoded and generalizes across types of depiction in early and high-level visual cortex

Based on previous findings (Haxby et al., 2001; Spiridon & Kanwisher, 2002; Walther et al., 2011), we hypothesized that information about the category of a presented object is represented in early and high-level visual cortex for natural images as well as for line drawings and that this information generalizes across levels of visual abstraction. To test this hypothesis, we trained and tested SVM classifiers on the fMRI data to decode the category of a presented object for each ROI and for every type of depiction separately. We focused on EVC and LOC as proxies for early and high-level visual processing, respectively. In addition, we explored the region pIPS in the dorsal stream since a growing body of evidence supports an important role of regions in the dorsal visual pathway for object recognition (see Freud et al. (2016) and Ayzenberg & Behrmann (2022) for a review) and has shown a selectivity for object format in these regions (Freud et al. 2018; Snow et al. 2011).

The category decoding results for EVC, LOC, and pIPS are shown in Fig. 2b. Category information could be decoded with accuracies significantly above chance from the voxel activity patterns from EVC, LOC, as well as pIPS for all types of depiction (*p*<0.001, sign-permutation test). When directly comparing decoding accuracies between types of depiction within an ROI, we found that there were no significant differences between any of the types of depiction in either EVC (all *p*>0.205, sign-permutation test, FDR-corrected), LOC (all *p*>0.083, sign-permutation test, FDR-corrected) or pIPS (all *p*>0.364, sign-permutation test, FDR-corrected). Finally, decoding accuracies for all types of depiction were higher in EVC than in both LOC and pIPS (all *p*<0.001, sign-permutation test, FDR-corrected) and higher in LOC than in pIPS (all *p*<0.001, sign-permutation test, FDR-corrected), which is expected given the strong visual differences between object categories in a given type of depiction. To control that these differences in decoding accuracies between ROIs are not simply driven by a larger number of voxels for any of the ROIs, we carried out the same decoding analysis after equating the number of voxels included in all ROI masks by randomly subsampling the bigger ROI masks. This control analysis led to comparable results, demonstrating that the differences between ROIs are not driven by a larger ROI size of any of the ROIs. In sum, this suggests that information about the category of a presented object is represented in early and high-level visual brain regions for all levels of visual abstraction.

To identify the degree to which category information generalizes between photos, drawings and sketches, we carried out cross-decoding. The rationale behind this approach is that if the classifier trained on data from one type of depiction (e.g. photos) can be used for data from another type of depiction (e.g. drawings), it is concluded that the underlying representational format is similar. We evaluated three different comparisons - photo-drawing, photo-sketch and drawing-sketch, resulting in three values for each ROI. We found significant cross-decoding accuracies between all types of depiction already in EVC but also in LOC and pIPS (all *p*<0.001, sign-permutation test; FDR-corrected, Fig. 2c). To evaluate the robustness of these findings, we also correlated the lower triangular parts of RDMs of different types of depiction with each other for each ROI separately. This led to qualitatively similar results, confirming the cross-decoding results.

Next, to further examine the degree of generalization between types of depiction, we asked if the decoding accuracies within types of depiction were different compared to the cross-decoding accuracies across types of depiction in each ROI. If these accuracies are not significantly different, this would indicate that the representation of object category is invariant to the type of depiction. If, however, the cross-decoding accuracies are lower than the decoding accuracies within type of depiction, this would indicate that the representation is tolerant but not invariant to the type of depiction (Hebart & Baker, 2018). The results for all comparisons are shown in Fig. 3. Accuracies across types of depiction were significantly lower than the corresponding accuracies within types of depiction for all comparisons in both EVC and LOC (all p<0.002, sign-permutation test, FDR-corrected). In pIPS, only the comparisons “Photo minus Photo-Sketch” and “Drawing minus Drawing-Sketch” reached significance (all p<0.003, sign-permutation test, FDR-corrected), while the other comparisons were only marginally significant (all p=0.051, sign-permutation test, FDR-corrected). The fact that these differences were less pronounced in pIPS might be explained by the overall smaller decoding accuracies in pIPS. Moreover, the differences in LOC and pIPS were significantly smaller than the ones in EVC, and the differences in pIPS were smaller than the ones in LOC (all p<0.004, sign-permutation test, FDR-corrected). These smaller effects in LOC and pIPS are consistent with the idea of gradually increasing tolerance to the type of depiction with depth of visual processing, yet, could also be explained by overall smaller decoding accuracies in LOC and pIPS. Overall, these results suggest that while the representation of object category in EVC, LOC and to some extent in pIPS is not invariant to the type of depiction, it exhibits tolerance to the type of depiction.

**Figure 3.**
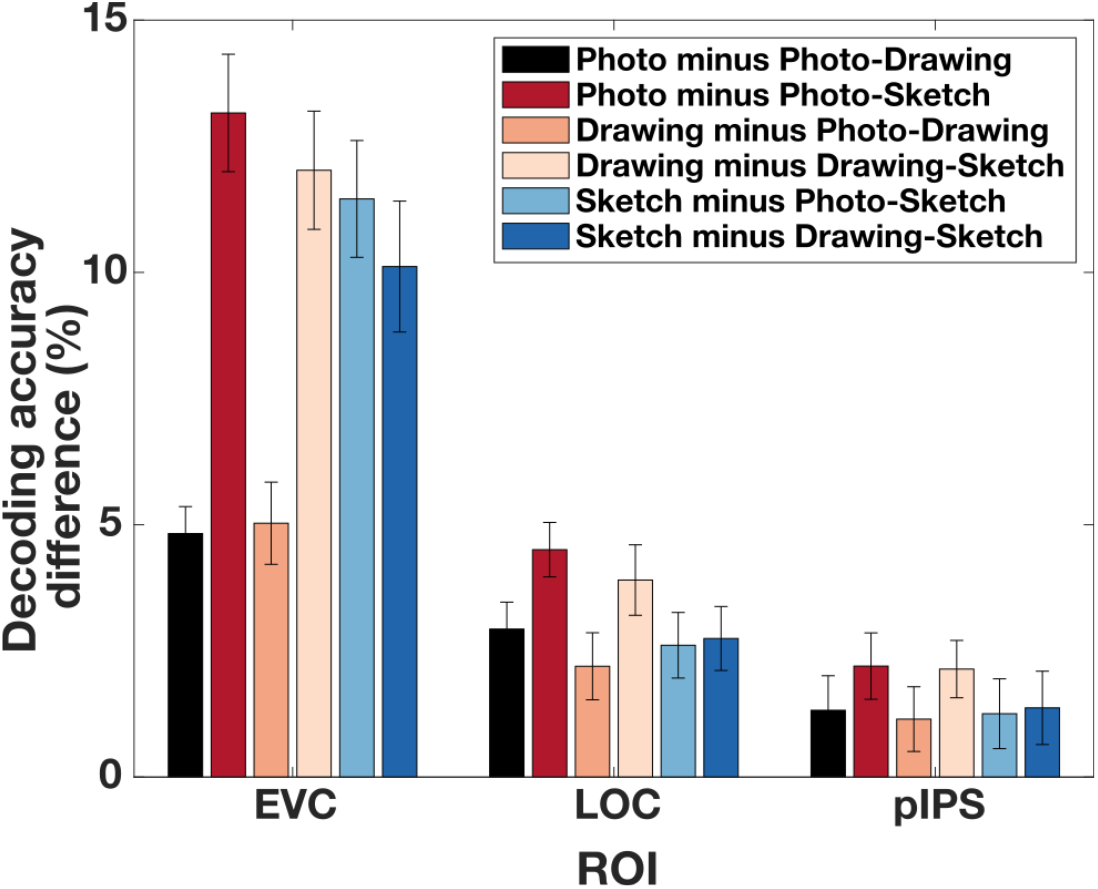
Tolerance for the type of depiction in low- and high-level visual cortex. Decoding accuracies within type of depiction were significantly higher than the decoding accuracies across types of depiction in EVC and LOC but only to some extent in pIPS. These differences were smaller for LOC and pIPS compared to EVC and smaller in pIPS than in LOC. Error bars reflect the standard error of the mean across participants.

Together, these findings corroborate earlier studies showing that category information can be decoded and is similarly represented in early and high-level visual cortex for natural object images and abstract drawings.

### 5.3. Large parts of occipital and ventral temporal cortex conjointly carry object category information which generalizes across levels of visual abstraction

While the results from the ROI analyses suggest a shared representational format of object category information across types of depiction in these ROIs, they leave open the spatial extent of this shared representation beyond these ROIs. To identify where category information is reflected in the brain across levels of visual abstraction and where it generalizes between types of depiction, we carried out a spatially unbiased searchlight analysis, iterating the decoding procedure over all possible searchlight locations in the brain.

The searchlight maps for decoding within types of depiction are shown in Fig. 4a. We found significant accuracies across large parts of occipital, ventral-temporal, and to some extent also posterior parietal cortex (*p*<0.05, cluster-based permutation test), with a strong overlap in the significance maps across types of depiction. Yet, significant voxels for photos extended more into anterior parts of ventral-temporal cortex than for drawings and sketches. To quantify the overlap between types of depiction, we conducted a conjunction analysis based on the intersection between all voxels that were significant for all three types of depiction (Nichols et al., 2005). The resulting conjunction map shows where category information was conjointly found across levels of visual abstraction (Fig. 4a). Confirming our initial observation, the conjunction map covered large parts of the occipital and ventral-temporal cortex, as well as a part of posterior parietal cortex. Beyond these similarities, no significant differences in decoding accuracies were found between different types of depictions (all *p*>0.05, cluster-based permutation test).

**Figure 4.**
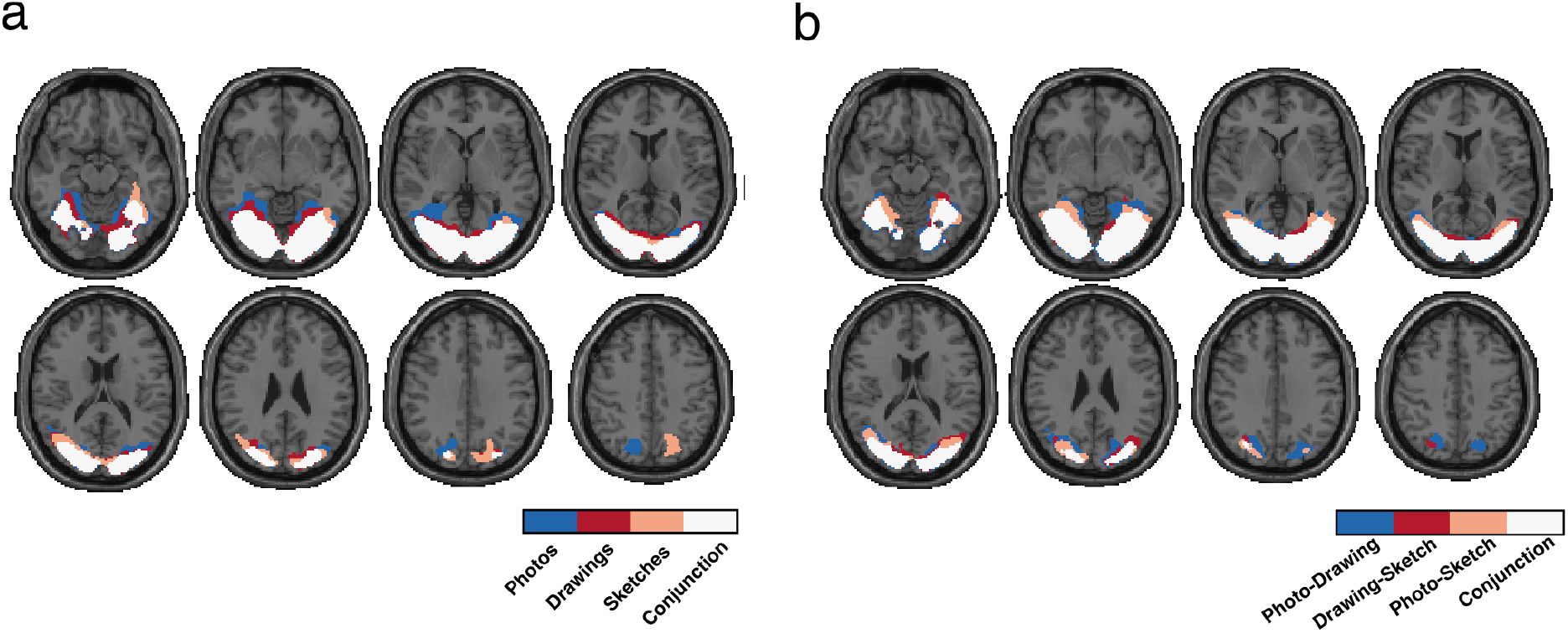
Representation and generalization of category information at different levels of visual abstraction across the whole brain. **a) Searchlight significance maps of the decoding of object category across levels of visual abstraction.** The color-coded masks indicate individually significant voxels for the decoding accuracies for each type of depiction separately. The conjunction map (color-coded in white) indicates conjointly significant voxels for all types of depiction. While significant areas for photos spanned more anterior parts of ventral temporal cortex than the ones for drawings and sketches, overall we found large parts of occipital and ventral-temporal and a part of posterior parietal cortex that conjointly reflected category information regardless of the level of visual abstraction. **b) Significance maps of the cross-decoding of object category between types of depiction.** Searchlight cross-decoding across the whole brain resulted in significant accuracies between all types of depiction in large parts of occipital and ventral-temporal cortex as well as a part of posterior parietal cortex. The conjunction map revealed a broad overlap in the locus of the generalizable information between all types of depiction.

The results for the searchlight cross-decoding between different types of depiction can be seen in Fig. 4b. We found significant cross-decoding accuracies between all types of depiction in large regions in occipital and ventral-temporal cortex, and to a smaller extent in posterior parietal cortex (*p*<0.05, cluster-based permutation test). The conjunction map for all three types of comparisons showed a broad overlap for all three comparisons mirroring the results from the within-type decoding.

In sum, this suggests that - beyond localized regions in early and high-level visual cortex - a large part of the ventral visual stream as well as parts of the dorsal visual stream reflect information about the object in a format that can be generalized across different levels of visual abstraction of the image.

### 5.4. Category information can be decoded rapidly from MEG activity patterns and generalizes early across types of depiction

Having established where category information can be decoded and where it generalizes across types of depiction, we investigated when information about the category of a presented object can be read out and when this information generalizes across types of depiction. Assuming that drawings and natural object images are similarly processed from early on in the visual system (Sayim & Cavanagh, 2011), we expected (1) that object category information should emerge with similar temporal dynamics for all types of depiction and (2) that category information should generalize early. In contrast, if additional processing is required to resolve the abstract visual information in drawings, we expected delayed emergence of category information for drawings and sketches in comparison to photos and generalization of category information only late in time. To distinguish between these alternatives, we trained and tested SVM classifiers either on the MEG channel patterns from the same or different types of depiction to decode the category of a presented object for each time point analogous to the fMRI decoding procedure.

The results of the temporally-resolved MEG decoding analyses within photos, drawings and sketches are shown in Figure 5b. Irrespective of the type of depiction, there was a rapid early rise in decoding accuracy, followed by a steady decline that continued into the end of the trial and that remained significant for all three levels of depiction (*p*<0.05, cluster-based permutation test). Overall, time courses were very similar for the three conditions, peaking at 100ms for all conditions (photo peak 95% confidence interval (CI) = [99.91ms 100.09ms], drawing peak CI = [98.61ms 101.45ms], sketch peak CI = [86.15ms 105.49ms]), with no significant differences between peak latencies (all *p*>0.05, based on bootstrap CI). A two one-sided tests procedure (TOST) testing for statistical equivalence of the peak latencies revealed significant results for all comparisons (Photo vs. Drawing, *MD*=-0.5ms, *p*<0.001; Photo vs. Sketch, *MD*=5ms, *p*=0.004; Drawing vs. Sketch, *MD*=5.5ms, *p*=0.007, FDR-corrected). Despite these similarities, the overall accuracy for the three conditions was different, as highlighted in the difference time courses (Fig 5c). There were significantly higher decoding accuracies for photos than for both drawings and sketches and significantly higher decoding accuracies for drawings than for sketches (all p<0.05, cluster-based permutation test). These differences suggest that there was a gradual decrease in the strength of the representation of category information with an increasing level of visual abstraction potentially related to the additional visual information (e.g. color, texture) contained in photos and drawings.

**Figure 5.**
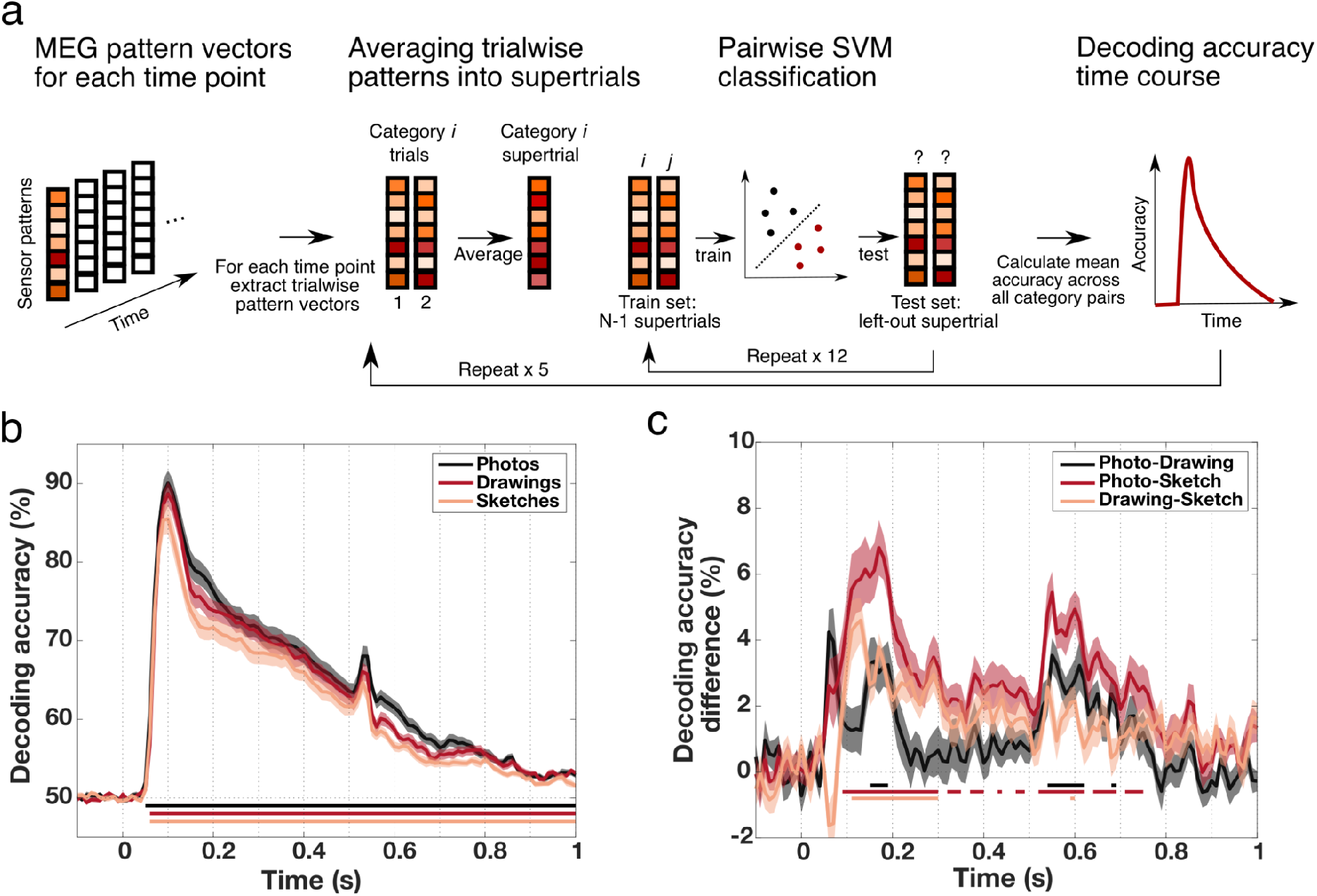
MEG-based category information resolved in time across levels of visual abstraction. **a) Temporally resolved decoding procedure.** For each time point, an SVM classifier was applied to MEG channel pattern supertrials in a repeated leave-one-out cross-validation scheme for all object categories *i* and *j.* The resulting decoding accuracies were then averaged over all possible object pairs and all repetitions for every time point. This yielded decoding accuracy time-courses for every participant and every type of depiction separately. **b) Temporally resolved decoding accuracies across levels of visual abstraction.** For all types of depiction, category information emerged rapidly after stimulus presentation, peaking at 100ms and gradually declining afterwards. The decline was interrupted by a small increase in accuracies shortly after stimulus offset around 530ms. **c) Differences in decoding accuracies between types of depiction.** When directly comparing decoding accuracies between types of depiction across time we found that accuracies were significantly higher for photos compared to both drawings and sketches. Additionally, drawing accuracies were higher than sketch accuracies. Shaded areas represent the standard error of the mean across participants for each time point. Colored lines below the accuracy plots indicate significant time points (*p*<0.05, cluster-based permutation test).

The cross-decoding time courses, which are depicted in Fig. 6a, showed a similar pattern for all comparisons, with a sharp increase shortly after stimulus presentation leading up to a peak after which accuracies declined slowly and remained significant for all three comparisons up until the end of the trial (*p*<0.05 cluster-based permutation test). Accuracies for all three comparisons peaked at 100ms (photo-drawing 95% peak CI = [93.40ms 104.78ms], photo-sketch CI = [85.54ms 105.21ms], drawing-sketch CI = [86.18ms 105.58ms]) with no significant differences between any of the peak latencies (all *p*>0.05, based on bootstrap CI). Testing for equivalence of the peak latencies revealed significant results for the comparison of photo-drawing and drawing-sketch peaks (*p*=0.008, TOST, FDR-corrected) but non-significant results for the other two comparisons (both *p*=0.24, TOST, FDR-corrected). Please note that an analysis correlating the lower triangular parts of RDMs of different types of depiction with each other for each time point led to comparable results, corroborating these findings.

**Figure 6.**
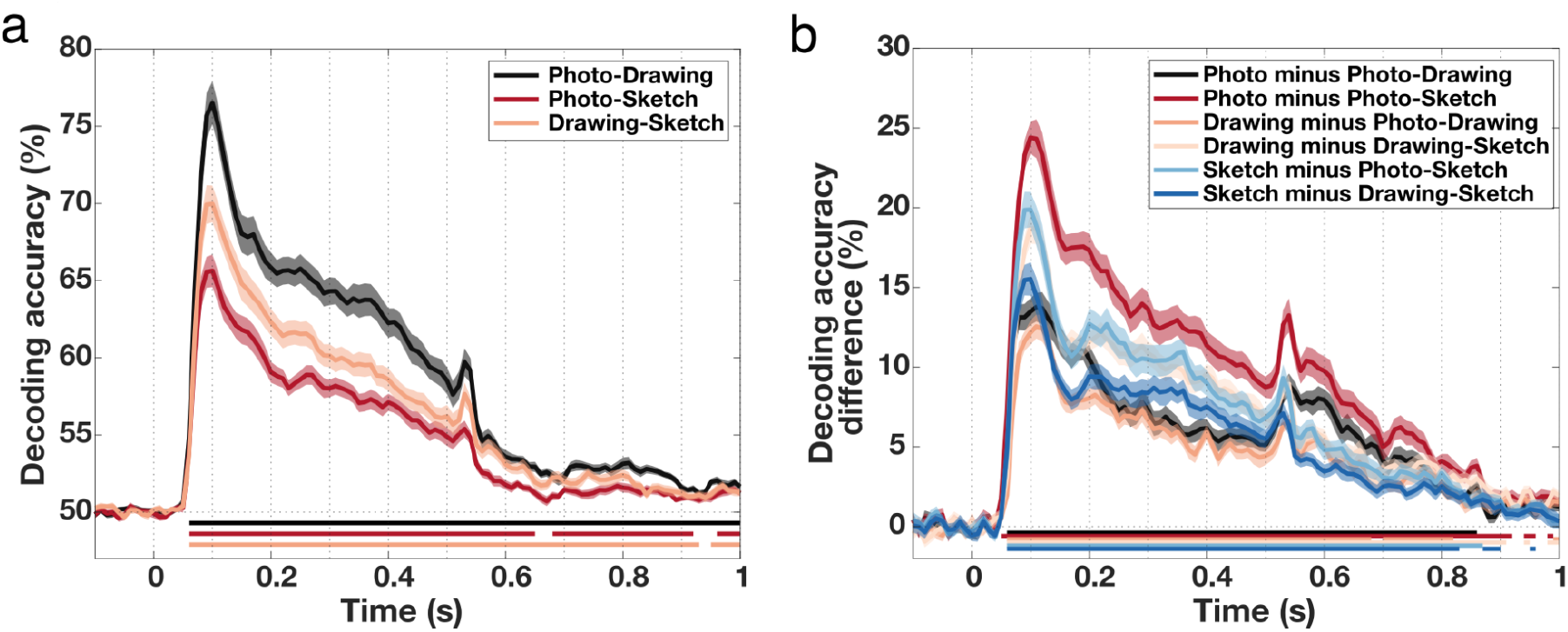
Generalization of object category information between types of depiction across time. **a) Cross-decoding between types of depiction across time.** Between all types of depiction, we found high cross-decoding accuracies based on the MEG data already early on, peaking around 100ms, and remaining high until shortly after the offset of the stimulus. **b) Differences between decoding within and across types of depiction across time.** Beginning early, peaking around 100ms, and remaining throughout most of the trial, decoding accuracies within type of depiction were significantly higher than decoding accuracies across types of depiction. These differences declined after the peak. Shaded areas reflect the standard error of the mean across participants for each time point. Colored lines below the accuracy plots indicate significant time points (*p*<0.05, cluster-based permutation test).

Moreover, to further assess the generalization between types of depiction, we compared decoding accuracies within types of depiction with accuracies across types of depiction in a time-resolved fashion. We found significantly higher decoding accuracies within types of depiction for all comparisons, which remained significant throughout most of the trial (p<0.05, cluster-based permutation test; Fig. 6b). Differences increased rapidly, peaked early around 100ms, and declined afterwards. In line with the results from the fMRI data, these results suggest that the representation of object category is tolerant rather than invariant to the type of depiction.

Together, these results show that object category can be decoded from stimulus evoked brain activity for natural images and drawings regardless of the level of visual abstraction, with similar temporal dynamics but a larger effect for natural object images which decreased across levels of visual abstraction. Furthermore, object category information generalized strongly across all levels of visual abstraction beginning already in early stages of visual processing and persisting into late stages of visual processing. This suggests that recognition of drawings and natural object images share strong similarities from early on in visual processing.

### 5.5. Comparable generalization of category information across time for all levels of visual abstraction

The temporally-resolved decoding analyses suggest that object category information emerges similarly fast for all types of depiction and generalizes early across depiction types. Yet, there might be differences in the dynamics and the stability of the representations between levels of visual abstraction. Such differences in the temporal dynamics between types of depiction would indicate differences in the underlying neural mechanisms for recognition of natural object images and line drawings. To investigate how the representation of category information for photos, drawings and sketches generalizes across time, we used temporal generalization analysis (King & Dehaene, 2014; Meyers et al., 2008), training a classifier on one time point and testing on all other time points for every type of depiction separately.

The resulting time × time generalization matrices for photos, drawings and sketches are shown in Fig. 7a. We found a similar pattern for all three types of depiction with strong generalization of the representation of category information beginning shortly after stimulus onset and continuing across the whole trial period (*p*<0.05, cluster-based permutation test). For all types of depiction there was a strong on-diagonal pattern from 50ms to ~200ms with comparatively weak off-diagonal accuracies early on. Later on, there was a stronger off-diagonal component after ~200ms until ~500ms. The overall pattern observed in the temporal generalization matrices was qualitatively similar across types of depiction indicating that the representation of the category of a presented object underwent comparable representational transformations in time for all types of depiction.

**Figure 7.**
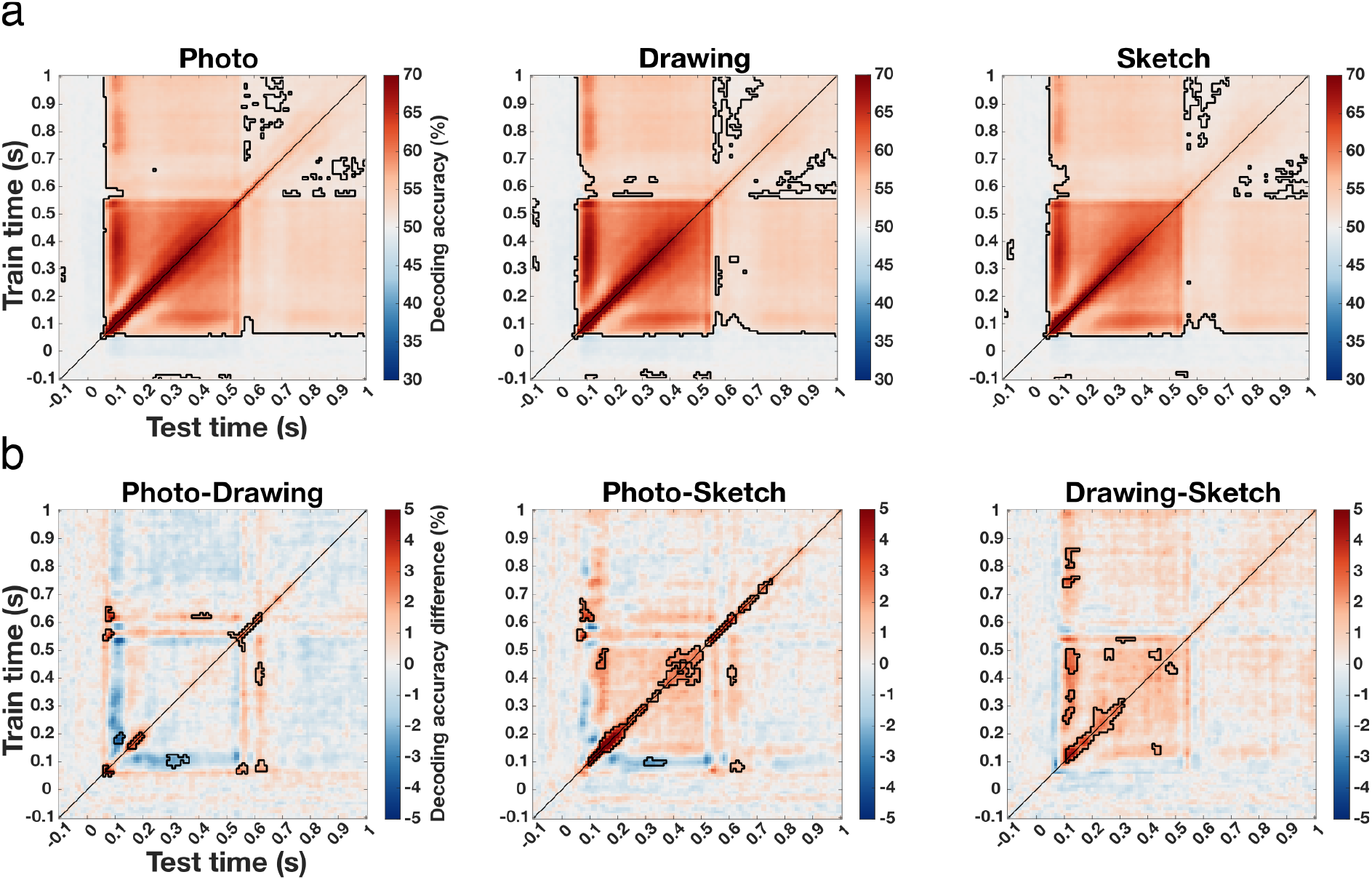
Generalization of the representation of category information across time for all types of depiction. **a) Temporal generalization matrices for the three types of depiction.** For all three types of depiction, we found strong generalization across time covering large parts of the trial. The pattern for the three types of depiction was qualitatively similar with a strong early on-diagonal component followed by a later component which showed additional strong off-diagonal elements. **b) Differences in temporal generalization between types of depiction.** The direct comparison of the temporal generalization between types of depiction revealed that there were differences in the strength of generalization. These differences between photo and sketches as well as drawings and sketches were most pronounced for on-diagonal elements. In addition, we found differences between photos and drawings which were less pronounced and without a clear pattern. Significant time points are indicated by the outlined areas (*p*<0.05, cluster-based permutation test).

The direct comparison of the pattern of generalization between photos, drawings and sketches, shown in Fig. 7b, revealed significant differences between all depiction types (*p*<0.05, cluster-based permutation test). Accuracies for photos were overall higher than for sketches, with the strongest differences spanning on-diagonal elements. Moreover, there were significantly higher decoding accuracies for drawings than for sketches. The strongest differences again mostly covered on-diagonal elements, yet with some distributed off-diagonal differences. For the comparison of photos and drawings the differences were less strong and did not show a clear pattern as for the other comparisons. Significant differences were more distributed with higher values for photos mostly on the diagonal and also some off-diagonal elements showing higher values for drawings.

In sum, these results demonstrate similarities in the overall pattern of generalization of category information across time but also differences in the strength of generalization. These differences were strongest for on-diagonal elements for the photo-sketch and drawing-sketch comparison, suggesting differences in the overall representation of category information between types of depiction but less so for the generalization across time. Differences between photos and drawings were less pronounced and scattered, limiting a strong interpretation of these results.

### 5.6. Similarities and differences in the combined spatiotemporal dynamics of object recognition for different levels of visual abstraction

Our results so far suggest that there are broad commonalities in the spatial and temporal dynamics of the representation of object category across levels of visual abstraction. Further, object category information generalized strongly from early visual processing stages on. Yet, the temporally-resolved decoding results and temporal generalization results indicate that there were differences in the strength of representation between types of depiction while the spatially-resolved decoding results did not show such differences. Hence, the question remains where differences in the neural dynamics between photos, drawings and sketches arise and at what time they arise in a given region. To combine the temporal and spatial information from MEG and fMRI data, we used RSA-based MEG-fMRI fusion (Cichy et al., 2014; Cichy & Oliva, 2020; Hebart et al., 2018). We computed RDMs for each ROI for the fMRI data and for each time point for the MEG data and and correlated the lower triangular parts of the temporally-resolved and ROI-specific RDMs (see Fig. 8a for visualization of the approach). This way, we could ask in what ROI and at what point in time the representation of objects was similar, revealing the spatiotemporal dynamics of object processing for photos, drawing and sketches. For visualization purposes we also carried out the MEG-fMRI fusion analysis using a spatially unbiased searchlight approach (Cichy et al., 2016), iterating the RDM correlation across searchlights and timepoints. The resulting MEG-fMRI fusion movies are publicly available via https://osf.io/vsc6y/.

**Figure 8.**
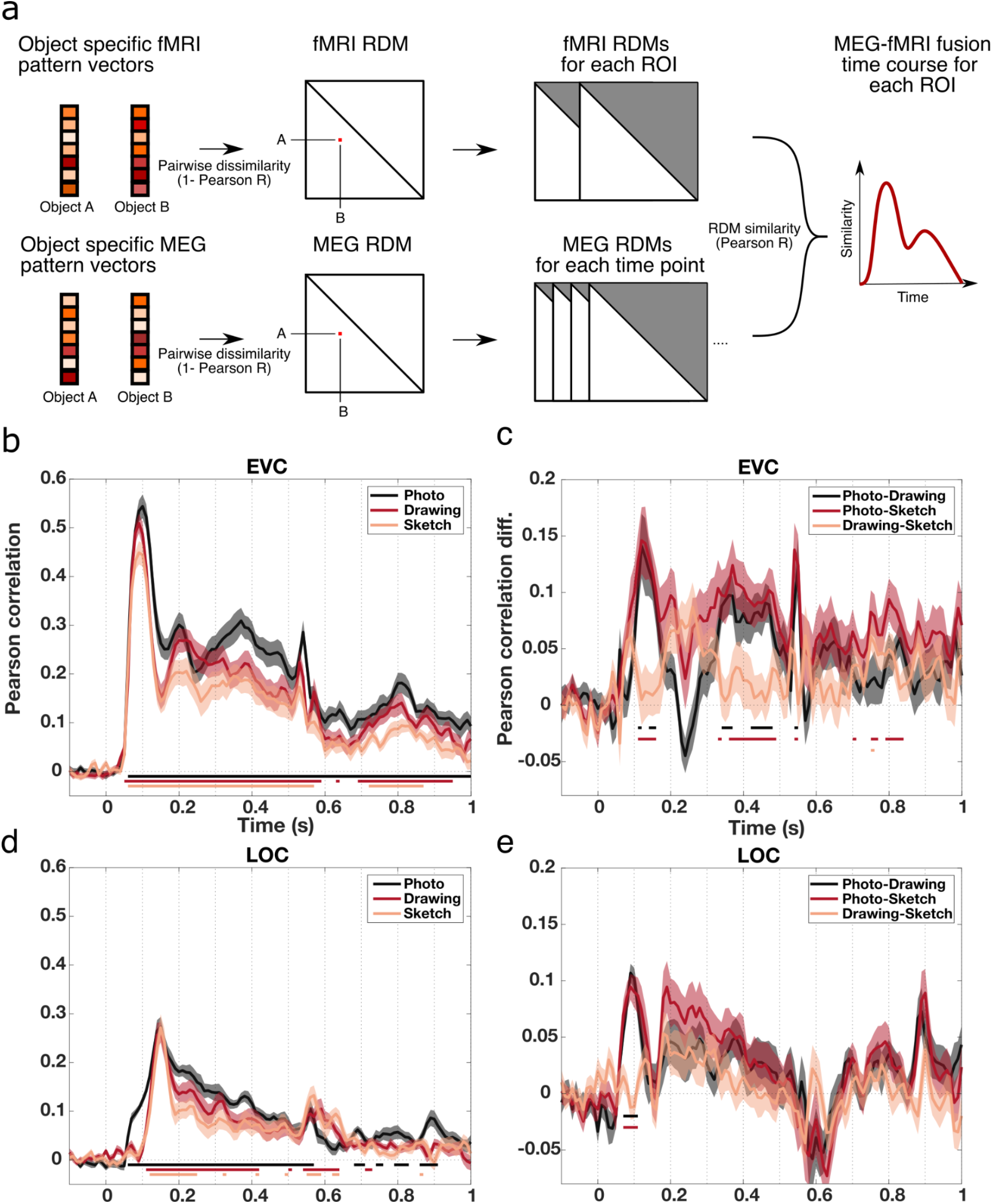
Spatiotemporal dynamics of object recognition for different levels of visual abstraction in EVC and LOC. **a) RSA-based MEG-fMRI fusion procedure.** For combining the spatial and temporal information from fMRI and MEG we first computed RDMs by calculating the pairwise dissimilarities (1-Pearson correlation) between all object-specific pattern vectors for every ROI or time point and every type of depiction separately. Then we correlated the lower triangular parts of the ROI-wise and timewise RDMs for each ROI and every type of depiction separately. This yielded MEG-fMRI fusion time courses for each ROI reflecting the spatiotemporal dynamics of object recognition for photos, drawings and sketches. **b) MEG-fMRI fusion time courses in EVC.** In EVC we found an early peak in MEG-fMRI correlation around 100ms for all types of depiction, with no differences in peak latencies. **c) MEG-fMRI fusion difference time courses between types of depiction in EVC.** Photos showed a stronger correlation than both drawings and sketches in EVC, particularly around 100ms to 200ms and around 300ms to 500ms after stimulus presentation. The differences between drawings and sketches in EVC were small but significant. **d) MEG-fMRI fusion time courses in LOC.** MEG-fMRI correlations in LOC peaked significantly later than in EVC around 150ms. There were no differences between peak latencies of different types of depiction. **e) MEG-fMRI fusion difference time courses between types of depiction in LOC.** In LOC there were no significant differences between drawings and sketches while photos showed a stronger correlation than both drawings and sketches particularly early on before 150ms. Shaded areas represent the standard error of the mean across participants for each time point. Colored lines below the accuracy plots indicate significant time points (*p*<0.05, cluster-based permutation test).

The fusion time courses for all types of depiction in EVC and LOC are shown in Fig. 8b and d respectively. In EVC we found an early increase in MEG-fMRI correlation for all types of depiction leading up to peaks, followed by a sharp decrease and another rise. After this second rise in correlation the MEG-fMRI correlations slowly decayed for drawings and sketches while for photos there was another increase. Finally, there was a last spike in correlation for all types of depiction around 500 to 540ms likely reflecting effects induced by the offset of the stimulus. Peak latencies for all types of depiction were found in the time from 90ms to 100ms (95% CI photo = [89.47ms 106.55ms], drawing = [83.32ms 95.29ms], sketch = [81.63ms 103.06ms]) with no significant differences between any types of depiction (all *p*>0.05, based on bootstrap CI of difference). Moreover, we tested for equivalence of the peak latencies which revealed non-significant results for all comparisons (all *p*>0.626, FDR-corrected). The comparison of fusion time courses between photos and both drawings and sketches in EVC, shown in Fig. 8c, revealed that there were significantly higher correlations for photos than for both drawings and sketches (*p*<0.05, cluster-based permutation test). These differences were strongest in the time around 100ms to 200ms and the time from ~300ms to ~500ms. Differences between drawings and sketches in EVC were only small but significant (*p*<0.05, cluster-based permutation test). In sum, object information regardless of the level of visual abstraction first peaked in EVC at around 100ms and then re-emerged later around 200ms, after which the representation slowly decayed for drawings and sketches, while for photos there was another late rise.

In LOC we found a rise in correlation for all types of depiction with peaks at 150ms for all three types of depiction (95% CI photo = [141.60ms 155.65ms], drawing = [136.50ms 156.39ms], sketch = [146.03ms 154.92ms]), significantly later than the peak latencies in EVC for all types of depictions (all *p*<0.05, based on bootstrap CI of difference; Fig. 8d). After these peaks, the correlation decayed up until the end of the trial only interrupted by a small rise shortly after the offset of the stimulus. There were no significant differences between peak latencies of different types of depiction in LOC (all *p*>0.05, based on CI of difference). Testing for statistical equivalence revealed non-significant results for all comparisons of peak latencies (all *p*=0.841, TOST, FDR-corrected). Furthermore, MEG-fMRI correlations were stronger for photos than for both drawings and sketches in LOC while there were no significant differences between drawings and sketches (*p*<0.05 cluster-based permutation test; Fig. 8e). Significant differences between photos and both drawings and sketches in LOC were mostly confined to early time points before 150ms.

Finally, we also explored the spatiotemporal dynamics of visual processing for photos, drawings and sketches in the region pIPS in the dorsal visual pathway. The MEG-fMRI correlation and MEG-fMRI correlation difference time courses for pIPS are shown in Fig. 9a and 9b, respectively. In pIPS, correlations increased up to a peak at 130ms for photos and drawings and at 150ms for sketches (95% CI photo = [57.15ms 207.54], drawing = [6.95ms 312.74ms], sketch = [91.27ms 204.49ms]), followed by a sharp decrease and a late rise in correlation after the offset of the stimulus. No significant differences between peak latencies were found (all *p*>0.05, based on CI of difference) and equivalence tests revealed non-significant results for all comparisons (all *p*=0.99, TOST, FDR-corrected). Please note that due to overall weaker effects in pIPS, the first peak in the time series could not be reliably detected using the whole trial period for detection. Therefore, we restricted the peak detection to the time period from the beginning of the trial up to the time the stimulus presentation ended (−100ms to 450ms). Moreover, we found significant differences between the correlations for all types of depiction (*p*<0.05 cluster-based permutation test, Fig. 9b). However, these differences were overall rather small and did not follow a clear pattern, limiting strong interpretation of these effects.

**Figure 9.**
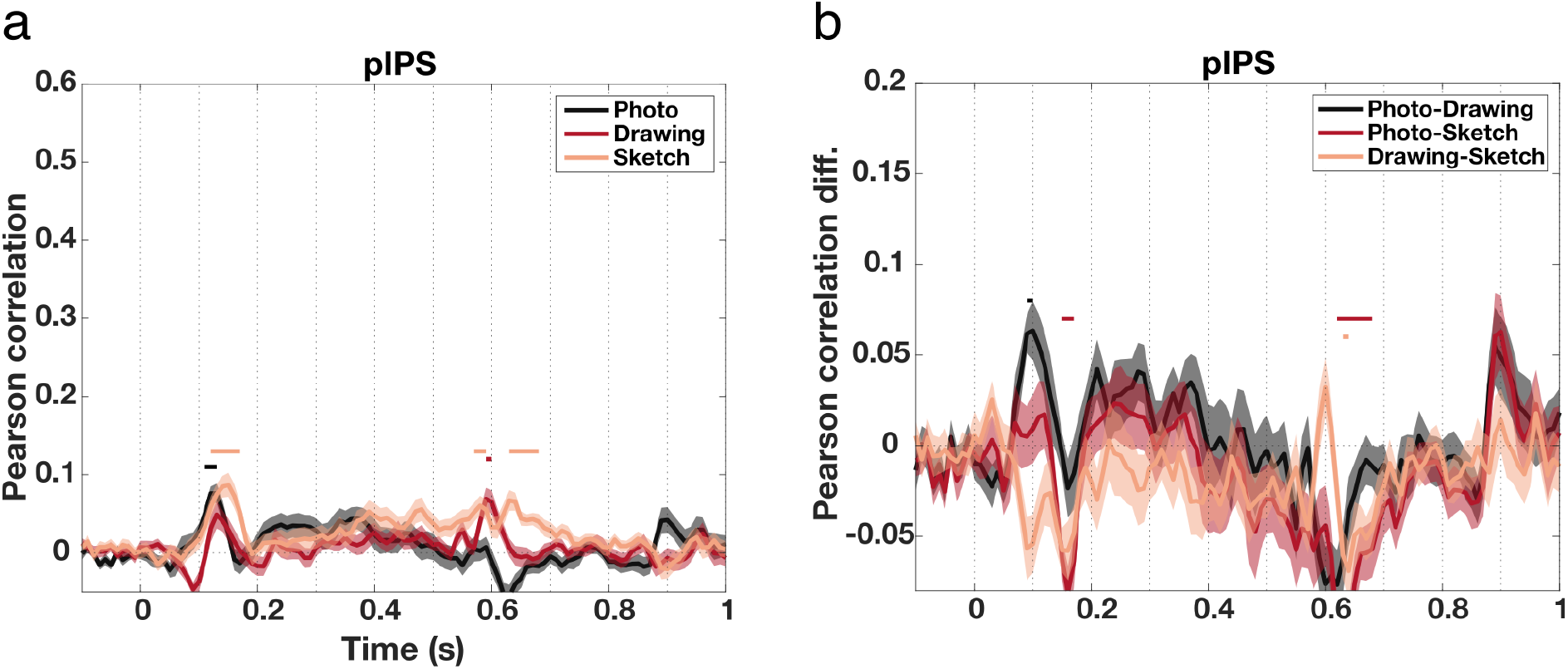
Spatiotemporal dynamics of object processing for different levels of visual abstraction in pIPS. **a) MEG-fMRI fusion time courses in pIPS.** For all types of depiction, the MEG-fMRI correlation increased early and peaked at 130ms for photos and drawings and for 150ms for sketches. There were no significant differences between peak latencies. **b) MEG-fMRI fusion differences in pIPS.** Differences in MEG-fMRI correlations between types of depiction in pIPS were small but significant. Overall these effects were rather unstable changing directionality over time. Shaded areas represent the standard error of the mean across participants for each time point. Colored lines above the accuracy plots indicate significant time points (*p*<0.05, cluster-based permutation test). The y-axes were scaled to be consistent with Figure 8b-e.

Taken together, the spatiotemporal dynamics of object recognition followed a comparable pattern across levels of visual abstraction. For all types of depiction, object information first peaked in EVC and later in LOC. This was followed by re-emergence of object information in EVC and a late phase of object processing with a sustained response in EVC. In addition, even in high-level visual regions in the dorsal visual pathway there was no difference in the emergence of object information between types of depiction. Despite these similarities, photos were distinctive in terms of the strength of representational similarity between fMRI and MEG data and showed both early and late differences in EVC as well as particularly early differences in LOC. In pIPS, differences were overall less pronounced and less stable, making the interpretation of these effects more challenging. In sum, these results indicate additional processing at multiple stages for photos in comparison to both drawings and sketches.

## 6. Discussion

In this study, we sought to identify the spatiotemporal neural dynamics underlying the processing of object drawings and to determine the similarities and differences to the processing of natural object images. Specifically, we used fMRI and MEG to distinguish between two alternative predictions: That photos, drawings, and sketches share the same representational format in both space and time, or that, alternatively, additional, potentially time-consuming processes would be required for the recognition of drawings and sketches. While these two predictions are not mutually exclusive, our findings only confirm the former prediction in four ways. First, we demonstrated that information about the category of a presented object could be read out from brain activity similarly fast and across large parts of the ventral visual stream as well as posterior parietal cortex, regardless of the type of depiction of the image. Second, the representation of object category information generalized beginning early in visual processing. Third, results from temporal generalization analyses suggest that there were qualitatively similar temporal dynamics for photos, drawings and sketches. Finally, the MEG-fMRI fusion results indicate that visual information processing follows similar stages, first peaking in EVC and then later in LOC for all types of depiction, with similar dynamics even in pIPS outside the ventral visual stream. In sum, this demonstrates that there are broad temporal and spatial commonalities in the neural dynamics as well as similar underlying representations for natural images and drawings from early on in visual processing.

In addition, we did not find evidence confirming the latter prediction proposing additional processing for drawings and sketches. Rather, our results suggest the opposite, that is, enhanced processing for photos at multiple stages. We found a gradual decline in the strength of category representations across levels of visual abstraction in the MEG data, with photos showing the strongest representation, followed by drawings and sketches. Moreover, the comparison of the spatiotemporal dynamics between types of depiction showed that photos exhibited a stronger representation both early and late in time in early visual brain regions, and exclusively early on in high-level visual cortex as compared to both drawings and sketches.

Collectively, our findings substantiate the hypothesis that line drawings resemble natural object images in terms of some core visual features (Fan et al., 2018), leading to a similar representation of drawings and natural images in the brain (Sayim & Cavanagh, 2011). Contrary to the hypothesis of additional processing for the recognition of line drawings, our results suggest that more in-depth processing is elicited by natural object images at multiple stages. Finally, these results indicate that the same neural mechanisms that support natural object recognition might also hold for drawings across different levels of visual abstraction.

Despite the abstraction of substantial amounts of visual information in line drawings, we found broad commonalities in the neural dynamics of object recognition for natural object images and line drawings. In combination with earlier findings (Haxby et al., 2001; Lowe et al., 2018; Spiridon & Kanwisher, 2002; Walther et al., 2011), these results provide evidence for the hypothesis that the information retained in line drawings serves as a basis for visual recognition, consistent with an edge-based account of recognition (Biederman & Ju, 1988). However, our results also show that object representations are stronger for photos compared to drawings or sketches. This is consistent with the theory on the role of surface information in object recognition (Tanaka et al., 2001) and empirical evidence (for a review see: Bramão et al., 2011) which propose that visual information only contained in natural images such as color and texture exerts influence on object recognition. Our findings substantiate this notion and suggest that while edge-based information in drawings might be sufficient to elicit qualitatively similar spatiotemporal representational dynamics as for natural images, surface information significantly contributes to object recognition at multiple processing stages.

Previous work has suggested a shared representational format for objects depicted as natural photographs or line drawings in early and high-level visual cortex (Haxby et al., 2001; Spiridon & Kanwisher, 2002), while for scenes such similarities have been shown to become stronger or to only arise late in the visual hierarchy (Lowe et al., 2018; Walther et al., 2011). Our results corroborate and extend earlier findings in object recognition by demonstrating that commonalities between natural object images and line drawings emerge early in time and early in the visual hierarchy. Yet, these results conflict to some part with previous work in scene recognition. This discrepancy might be explained by the fact that our stimulus set comprised a single exemplar instead of multiple exemplars per category (Lowe et al., 2018; Walther et al., 2011), which emphasizes low-level visual feature differences in decoding category information. Yet, it is possible that these partly conflicting findings point to a distinction in the representation and relevance of low-level visual features such as edges in object and scene recognition (Groen et al., 2017), which invites further exploration.

We demonstrated that object category information emerges similarly fast in the brain for abstract drawings as compared to color photographs. This suggests that object recognition can be resolved with the same amount of processing resources for different levels of visual abstraction of the image. This is consistent with previous computational work showing that representations for photographs and drawings at different levels of visual abstraction become highly similar when being processed in feedforward deep convolutional neural networks trained to categorize natural object images (Fan et al., 2018; Singer et al., 2022). While other work has demonstrated that additional recurrent processing is necessary for resolving degraded (Wyatte et al., 2012), occluded (Rajaei et al., 2019; Tang et al., 2018) or otherwise challenging images (Kar et al., 2019), our findings indicate that no additional mechanisms are needed for the robust recognition of abstract drawings. Future research could identify precisely in which cases visual recognition can be resolved with or without the need for additional processing which might serve as an important constraint for future efforts in modeling object recognition.

One difference in visual processing between photos and both drawings and sketches was found with MEG-fMRI fusion very early on in high-level visual cortex. LOC exhibited a faster rise of object representations for photos than for the other depiction types. While the source of this specific difference is unclear and not found for MEG and fMRI data alone, one possible explanation for this finding is the marked difference in the spatial frequency spectrum between drawings and photographs. While drawings and sketches contain mainly high spatial frequency information, photos additionally contain low spatial frequency information (Walther et al., 2011). This increased presence of low spatial frequency information may have contributed to an earlier rise of information related to rapid extraction of coarse information (Bar, 2003; Bar et al., 2006; Kveraga et al., 2007; Musel et al., 2014; Petras et al., 2019; Peyrin et al., 2010; Schyns & Oliva, 1994; Sugase et al., 1999). Future studies that carefully control spatial frequency in an image might reveal to what extent the spatiotemporal dynamics of object recognition are influenced by different spatial frequency patterns (Perfetto et al., 2020).

Previous work has shown that regions in the dorsal visual stream respond differently to real objects and images of objects (Freud et al. 2018; Snow et al. 2011). Therefore, we explored if a similar sensitivity for the type of depiction of an object (i.e. photo vs. drawing) can be observed in these regions. We found category information that could be generalized across types of depiction and no evidence for differences in the emergence of category representations between types of depiction in pIPS. However, we also found differences in the strength of the spatiotemporal dynamics of object recognition in pIPS, which may lend support to a differentiation of drawings and natural images in the dorsal stream. However, the effects in pIPS were overall smaller and less pronounced as compared to the results in occipito-temporal cortex, which might point to low SNR in this region, potentially constraining the conclusions that can be drawn from our results. Future investigations focusing specifically on the dorsal visual pathway could use experimental designs and imaging protocols tailored to these regions to be able to more clearly contribute to the growing evidence for the involvement of the dorsal visual pathway in object recognition (Freud et al. 2016; Ayzenberg & Behrmann 2022).

It should be noted that while we ensured that stimuli in different types of depiction are perceptually different, we did not explicitly control which details were included in the drawings and sketches and which not. Some details such as junctions and curvatures have been shown to crucially contribute to the recognizability of drawings (Walther et al., 2011; Walther & Shen, 2014). To rule out that differences in the representation of drawings and natural images can simply be explained by differences in the recognizability, we ensured that the participants were able to recognize all stimuli. Yet, the presence of some features in a drawing might determine if it is processed similarly as natural images in the visual system or not. While our results do not answer the question what features exactly allow for the recognition of drawings, they demonstrate that the features that are commonly retained lead to a similar representation of drawings and natural images in the brain. Disentangling what types of features contribute to the representations in the visual system is an overarching goal in visual neuroscience and ongoing efforts as well as future investigations might reveal distinct contributions of different features (Bankson et al., 2018; Groen et al., 2018).

### 6.1. Conclusion

In conclusion, our results show that the set of core visual features retained in line drawings is sufficient to elicit a processing cascade in the visual system that is remarkably similar to the one of natural images. This suggests that the same neural mechanisms that support natural object recognition might also hold for abstract line drawings. While we did not find any evidence for the involvement of additional processing for drawings, our findings indicate that visual information unique to natural images modulates visual processing at multiple stages. These results contribute to the understanding of how drawings convey meaning efficiently on the one hand and provide important insights into the neural mechanisms that underlie robust object recognition on the other hand.

## Acknowledgements

We thank Juliana Sarasty Velez, who created the drawings and sketches used in this study and provided all rights to the images.

